# The influence of the phylogenetic inference pipeline on murine antibody repertoire sequencing data following viral infection

**DOI:** 10.1101/2020.03.20.000521

**Authors:** Alexander Yermanos, Victor Greiff, Tanja Stadler, Annette Oxenius, Sai T. Reddy

**Affiliations:** Department of Biosystems Science and Engineering, ETH Zürich, Basel, Switzerland; Institute of Microbiology, ETH Zürich, Zürich, Switzerland; Department of Immunology, University of Oslo, Oslo, Norway

## Abstract

Understanding B cell evolution following vaccination or infection is crucial for instructing targeted immunotherapies when searching for potential therapeutic or virus-neutralizing antibodies. Antibody phylogenetics holds the potential to quantify both clonal selection and somatic hypermutation, two key players shaping B cell evolution. A wide range of bioinformatic pipelines and phylogenetic inference methods have been utilized on antibody repertoire sequencing datasets to delineate B cell evolution. Although the majority of B cell repertoire studies incorporate some aspect of antibody evolution, how the chosen computational methods affect the results is largely ignored. Therefore, we performed an extensive computational analysis on time-resolved antibody repertoire sequencing data to better characterize how commonly employed bioinformatic practices influence conclusions regarding antibody selection and evolution. Our findings reveal that different combinations of clonal lineage assignment strategies, phylogenetic inference methods, and biological sampling affect the inferred size, mutation rates, and topologies of B cell lineages in response to virus infection.

## Introduction

B cells are important for the clearance and neutralization of various infectious pathogens via interactions of their characteristic B cell receptor (BCR, or secreted version: antibodies). Sophisticated molecular mechanisms generate an ensemble of antibodies capable to interact with a vast number of foreign antigens. Part of this diversity is achieved through the initial somatic recombination of variable (V), diversity (D), and joining (J) germline segments, which together encode for the antigen-binding region of the antibody (Tonegawa, 1983). Further diversity is introduced via somatic hypermutation (SHM), in which mutations are selectively introduced into the antibody locus (Di Noia and Neuberger, 2007; Methot and Di Noia, 2017). These mutations can increase the binding strength to a particular antigen (referred to as affinity maturation) and occasionally confer a neutralizing phenotype against a particular pathogen. Some HIV-neutralizing antibodies, for example, require multiple rounds of SHM over long periods of time to induce neutralizing capabilities against a wide range of HIV strains (Wu *et al*., 2015; LaBranche *et al*., 2018; Landais and Moore, 2018; Klein *et al*., 2013). One can then compare these broadly neutralizing antibodies to the unmutated germline common ancestor (i.e. the original V-D-J recombined ancestor before SHM) to both identify critical mutations for pathogen neutralization and to discover new pathogen-binding variants (LaBranche *et al*., 2018).

The advent of high-throughput sequencing has allowed an unprecedented resolution with which B cell dynamics can be studied during infection. High-throughput immunoglobulin repertoire sequencing (Ig-seq) experiments aim to isolate and describe co-existing B cell populations within an organism, providing thousands to millions of sequences for subsequent analyses (Georgiou *et al*., 2014; Miho *et al*., 2018). While these sequencing reads contain the evolutionary histories of multiple, independent monoclonal antibody lineages, how exactly this multitude of data should be processed remains unclear. The general pipeline involves first assigning reads to a single antibody lineage based on some similarity criteria (e.g., germline genes, sequencing homology, CDR3 lengths), with the goal of clustering all sequences arising from a single V-D-J recombination event (Yermanos *et al*., 2018). The sequences for each clonal lineage are then used as input to a phylogenetic inference method, thereby producing a phylogenetic tree in which the recovered sequences are the tips and the root is the unmutated germline ancestor. These phylogenetic trees can then be compared using metrics such as branch lengths, number of sequences per tree, or tree imbalance, both within and across individuals. Previous work has demonstrated that phylogenetics can aid discovery of virus-specific antibodies (Zhu, Wu, *et al*., 2013). However, extracting relevant biological information from the mass of phylogenetic trees co-evolving within a single host, which we refer to as the “antibody forest”, remains less straightforward. While many different tools and bioinformatics practices exist, specifically for clonal lineage analysis (Ralph and Matsen IV, 2016; Briney *et al*., 2016; Schramm *et al*., 2016; Gupta *et al*., 2015, 2015), it remains unclear how custom pipelines of clonal lineage assignment, phylogenetic inference, and topological analysis impact our understanding and interpretation of B cell evolution.

In the case of assigning reads to a given V-D-J recombination event, some publications have relied solely upon V- and J-gene usage, whereas others have implemented requirements pertaining to complementary determining region 3 (CDR3) lengths or sequence homology (Bhiman *et al*., 2015; Bonsignori *et al*., 2016; Doria-Rose *et al*., 2014; Jackson *et al*., 2014; Soto *et al*., 2016; Stern *et al*., 2014). Furthermore, there exist several methods to construct phylogenetic trees, including distance-based metrics, maximum likelihood (ML), maximum parsimony (MP), and Bayesian inference (Stamatakis, 2006; Gascuel, 2006; Bouckaert *et al*., 2014). While there exist multiple simulation tools capable of exploring how inference method impacts the resulting phylogenetic trees derived from simulated B cell data (Yermanos *et al*., 2017; Davidsen and Matsen IV, 2018; Safonova *et al*., 2015; Weber *et al*., 2019), the extent of this influence on the evolutionary conclusions on experimental data remains largely unexplored. It remains unknown, for example, the extent of which the phylogenetic inference strategy impacts the biological conclusions pertaining to the evolutionary landscape across various infection cohorts.

Here, we explored how various bioinformatics pipelines shape evolutionary conclusions arising from Ig-Seq experiments. Using previously published time-resolved Ig-Seq experiments from blood-derived B cell and bone marrow (BM) PC (PC) repertoires (Kräutler *et al*., 2020), we analyzed the robustness of multiple conclusions based on phylogenetic analyses for three cohorts: uninfected mice, mice infected with low dose (acute) lymphocytic choriomeningitis virus (LCMV), and mice infected with high dose (chronic) LCMV. This unique experimental model provides a system in which a viral infection is either cleared within two weeks (in the case of the low-dose, acute infection) in a primarily CD8 T+ cell dependent manner or over the course of months in the case of mice receiving the high dose infection. It was expected that these three cohorts have distinct B cell evolutionary profiles given the sustained presence of virus and germinal centers in the case of chronic infected mice compared to the other two cohorts.

In particular, we investigated how (i) clonal lineage assignment strategies, (ii) phylogenetic reconstruction strategies, and (iii) biological sample strategies impacted results. First, we show that properties of the clonal lineages including size and number of trees highly depend upon the initial rooting strategies. Second, the resulting phylogenies depend upon the chosen inference method despite using the same B cell sequences as input. Furthermore, we demonstrated that leveraging known reference germline information improves Bayesian reconstruction of certain parameters, such as tree height. Finally, we observed that clonal lineage strategies and phylogenetic inference methods impact size and temporal resolution of public clonal lineages. Our findings both suggest a degree of caution when interpreting Ig-Seq data and highlight the importance of benchmarking pipelines commonly employed in systems immunology.

## Results

### Rooting strategy influences number, size and time-resolution of clonal lineages

Assigning sequences from bulk Ig-seq data is typically one of the first steps in reconstructing B cell clonal lineages (Yermanos et al., 2018). Although multiple tools specifically tailored to clonal lineage assignment now exist (Ralph and Matsen IV, 2016; Briney *et al*., 2016; Schramm *et al*., 2016; Safonova and Pevzner, 2019), a vast number of studies have performed some variation of first aligning the recovered antibody sequences to the reference germline sequences and subsequently clustering based on germline gene usage, edit distance sequence homology and/or CDR3 length (Stern *et al*., 2014; Doria-Rose *et al*., 2014; Bhiman *et al*., 2015; Tsioris *et al*., 2015). While different germline aligner tools have been previously compared (Marcou *et al*., 2018), how the aligner tools impact the repertoire fingerprint remains less characterized. We therefore compared the influence of various clonal lineage assignment pipelines following identical germline assignment of when analyzing recently published time-resolved Ig-Seq data (Kräutler *et al*., 2020). This sequencing data set consists of bulk heavy chain sequencing from 15 mice of the three different previously mentioned cohorts (uninfected, acute LCMV and chronic LCMV infection) and provides a unique opportunity to compare the influence of the bioinformatics processing pipeline across both multiple individuals and cohorts (Figure 1A). Ten mice were infected with either low- or high-dose LCMV (n_high-dose_=5, n_low-dose_=5), resulting in acute (resolved within two weeks) and chronic infections (resolved after months), respectively. Furthermore, five uninfected mice were included as a control. For each mouse (excluding one acute mouse), blood samples at 10, 20, 50, 60 and 70 days post infection (dpi) and bone marrow PCs (a subset of antibody secreting B cells) 70 dpi were collected, in all samples the heavy chain repertoires were subjected to Ig-seq (Kräutler *et al*., 2020).

**Figure 1.**
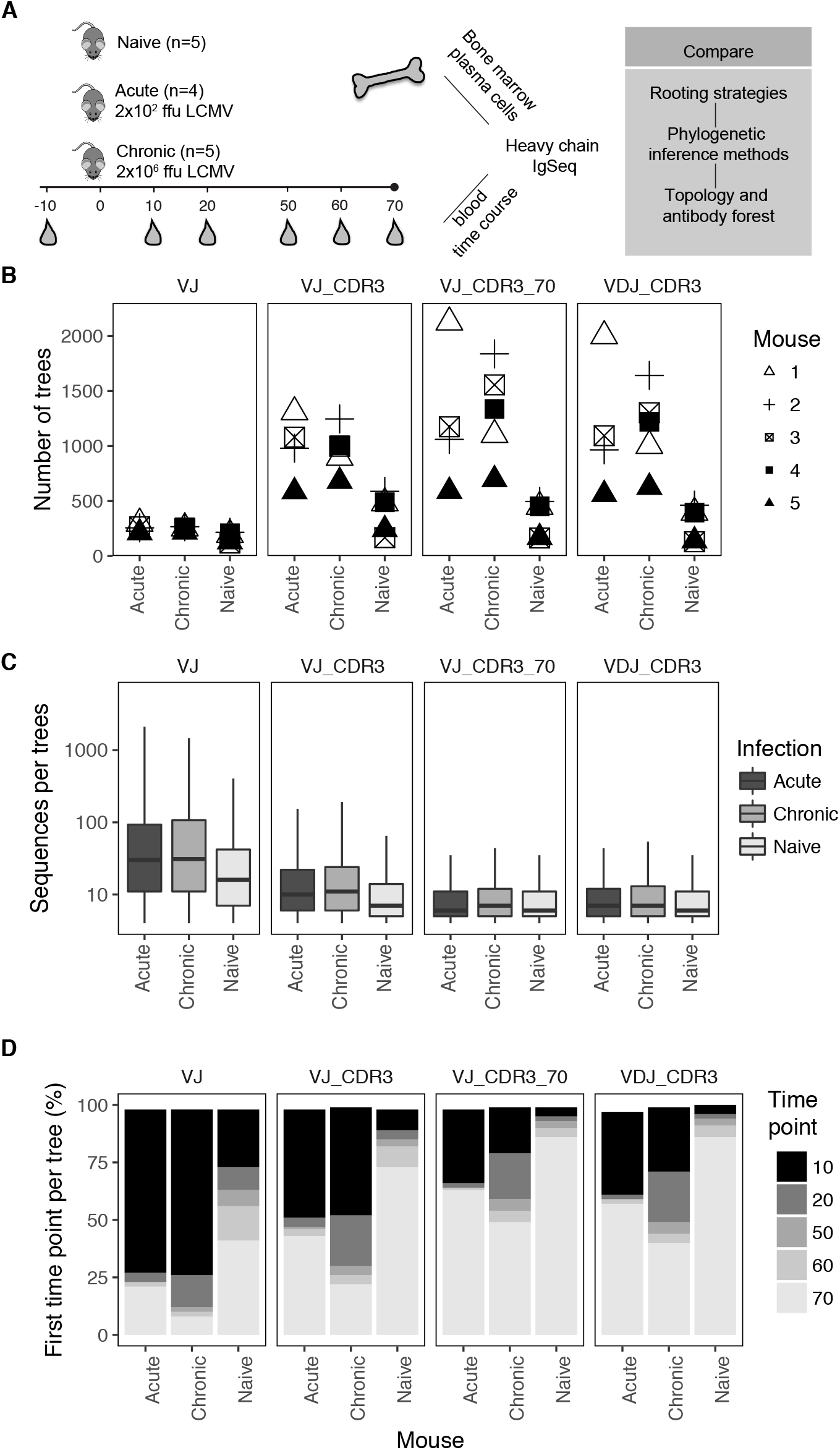
Rooting strategy influences phylogenetic fingerprint of different infection cohorts. (A) Schema depicting the experimental and computational pipeline. (B) The number of clonal lineages per mouse based on different clonal assignment strategies following germline assignment. (C) The average number of unique, full-length, sequences per clonal lineage following different clonal lineage assignment strategies. (D) The distribution of the first time points per tree average across all mice within a given cohort.

We first aligned the sequencing reads for each repertoire to the murine reference germline segments and subsequently defined clonotypes by the full-length VDJ region using MiXCR (Bolotin *et al*., 2015). After grouping all unique full-length VDJ nucleotide sequences for each mouse, we first asked how the quantity of clonal lineages for each mouse was impacted by clonal lineage assignment. We thereby assigned full-length VDJ sequences to a given clonal lineage if they shared the following characteristics: i) identical V- and J-gene germline segments (VJ), ii) identical V- and J-gene combination with the same CDR3 length (VJ_CDR3), iii) identical V- and J-gene combination with both the same CDR3 length and at least 70% CDR3 edit distance homology (VJ_CDR3_70), iv) and identical V-, D- and J-gene assignment with the same CDR3 length (VDJ_CDR3) (Figures 1B, S1A). Enumerating the number of clonal lineages per mouse revealed major discrepancies when comparing the size of the antibody forest for each of the aforementioned rooting methods (Figure 1B). Indeed, some rooting strategies, such as grouping sequences by V- and J-genes greatly reduced the number of clonal lineages for each mouse (Figure 1B, S1A). Other more stringent strategies to assign sequences to a given clonal family increased the number of clonal lineages as one may expect (Figure 1B). For all strategies, excluding grouping solely by V- and J-gene usage, the trends for both individual mice and cohort remained consistent despite the fact that the absolute number of clonal lineages varied across the different parameter settings (Figure 1B, S1A). This involved all mice infected with LCMV having more clonal lineages than the uninfected counterpart. This pattern was similarly observed when looking at the number of sequences per clonal lineage across each cohort (Figure 1C, S1B). We then quantified how the rooting strategy impacted the earliest time point observed within each clonal family, indicative of the time a given B cell was recruited into the immune response. Our analysis revealed that across all conditions the chronic cohort revealed a large proportion (at least 10%) of lineages that were first sampled at 20 dpi. While this continued recruitment of new clonal lineages during chronic viral infection was seen across all rooting strategies, the pipeline largely determined whether the majority of trees were seen already at 10 dpi or first at 70 dpi (Figure 1D). Finally, we asked how the number of time points per clonal lineage was impacted by the lineage assignment strategy. Similar to the earlier seen time point, the trend that chronically infected mice had more unique time points represented in each clonal lineage remained constant (Figure S2). Together, these results indicate that the number and size of clonal lineages is influenced by the rooting strategy, but after a certain threshold of stringency, for example, by forcing lineages to share identical CDR3 length or a high degree of homology, the trend across cohorts remains relatively stable. However, the temporal resolution (i.e. number of time points and earliest time point) of the clonal lineages produced partially contradicting conclusions, pending on imposing a homology requirement upon sharing V- and J-genes plus CDR3 length or not.

### Phylogenetic inference strategy alters tree topology

After observing the variability introduced solely due to the clonal lineage assignment strategy, we next questioned whether this noise was compounded by the phylogenetic inference method. Multiple inference methods and tools have been applied to Ig-seq data, including neighbor joining, maximum parsimony, maximum likelihood and Bayesian analysis (Jackson *et al*., 2014; Stern *et al*., 2014; Wu *et al*., 2015; Zhou *et al*., 2013; de Bourcy *et al*., 2017). We initially quantified whether two different inference methods resulted in identical tree topologies for the four aforementioned rooting strategies. Surprisingly, there was only minor congruence between tree topologies after inputting identical sequences into either neighbor joining or maximum likelihood pipelines (Figures 2A-C). For example, two distinct topologies were produced from identical sequences for a randomly selected clonal lineage from a chronically infected mouse (Figure 2A). Indeed, this phenomenon of divergent topologies despite identical input sequences was observed for all rooting strategies, with the V-J rooting strategy showing the highest divergence (Figure 2B). The more stringent methods requiring either identical CDR3 length, D-gene alignment, or a sequence homology threshold still resulted in merely ~30% of trees possessing identical topologies (ignoring branch length differences) (Figure 2B). Next, we questioned how similar the topology structure was for lineages containing identical sequences yet produced using different inference methods. For this, we calculated the normalized Robinson-Foulds (RF) distance, a commonly used metric that quantifies the number of partitions unique to each tree (RF distance of 0 = identical trees) (Robinson and Foulds, 1981). Similar to the percent of identical trees, the normalized RF distance revealed that the rooting method dramatically impacts how similar the topologies are despite identical input sequences, with the two most stringent rooting methods (VJ_CDR3_70 and VDJ_CDR3) possessing the lowest RF distances.

**Figure 2.**
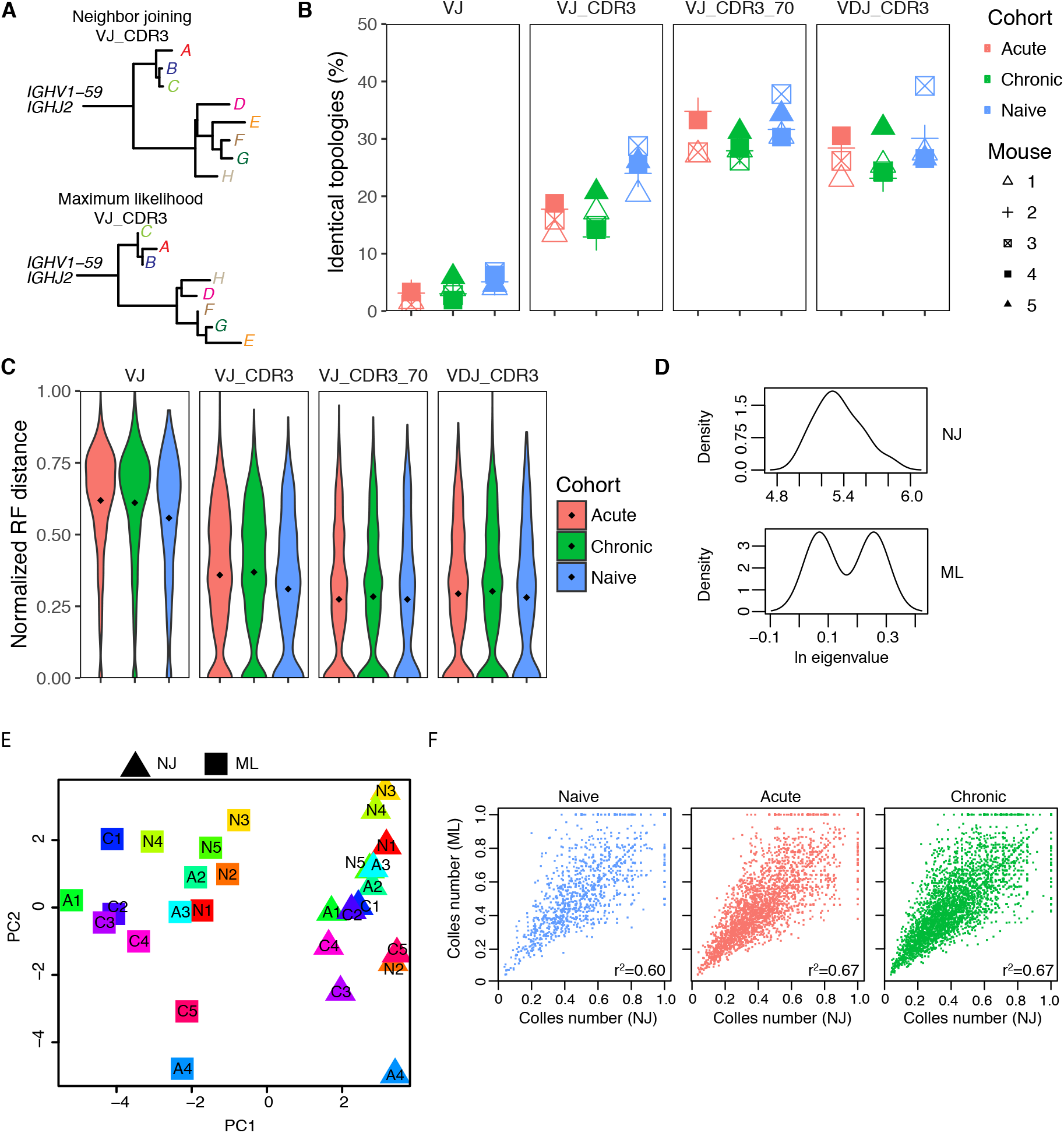
Clonal lineage assignment strategy and phylogenetic inference method alter evolutionary fingerprint across infection cohorts. (A) Representative phylogenetic trees inferred with identical sequences as input from a chronically infected mouse. (B) The percent of identical trees inferred by both maximumlikelihood inference (ML) and neighbor-joining (NJ) algorithms separate by moues and cohort. (C) The Robinson-Foulds distance normalized by the number of sequences quantifying the distance between trees containing identical input antibody sequences but inferred using either ML or NJ methods. Circles indicate the mean RF distance for each cohort. (D) Example Laplacian spectra of two different phylogenetic trees inferred with identical input sequences from A. (E) Principal component analysis based on the output from Laplacian for all trees across all mice using the VJ_CDR3 rooting strategy. Color indicates mouse and shape indicates infection cohort. Numbers correspond to individual mice within each cohort (N1=naïve mouse 1, A1=acute mouse 1, C1=chronic mouse 1). (F) Spearman correlation of the Colless number quantifying tree imbalance for each tree produced from identical sequences using either ML or NJ inference pipelines.

After observing such low agreement in tree topologies inferred using different methods for each clonal lineage, we questioned whether these topological differences alter metrics summarizing the trees. We first calculated the Laplacian spectra, a recently described metric that infers phylogenetic properties based on the underlying graph structure (Lewitus and Morlon, 2016), for all trees under the different rooting conditions. Plotting the spectral density from the exemplary trees (Figure 2A) revealed that, indeed, dramatic visual differences were distinguishable between the two phylogenetic inference methods (Figure 2D). To quantify this across mice and cohorts, we calculated the mean of multiple metrics describing the Laplacian spectra, such as the asymmetry, skewness, peakedness and the number of modalities (Lewitus and Morlon, 2016) and performed principal component analysis. A clear separation between the two inference methods was observed (Figure 2E). While the mouse-specific trend was somewhat constant for some mice (e.g. location of mouse A4 relative to the other mice in both conditions), for others the relative position dramatically shifted (e.g. C5 and N2 were much closer together for NJ trees and more distant for ML trees) (Figure 2E). We finally asked if other metrics that quantify tree imbalance produced similar discrepancies between the NJ and ML inference pipelines. To this end, we quantified the Colless number, a metric describing the balance of all internal nodes within a clonal lineage (Colless, 1982), for each NJ and ML tree across the three cohorts. Overall, the Colless number for trees containing identical sequences and potentially different topologies was similarly correlated across all three cohorts (r^2^=~0.6) (Figure 2F). Interestingly, those trees with higher Colless numbers showed more variability between NJ and ML methods, with some trees completely imbalanced (normalized Colless number = 1) in one method and not in the other. Taken together, our data suggest that the phylogenetic inference method influences tree topology and biological conclusions.

### Incorporating reference germline into Bayesian phylogenetics

In order to produce a rooted tree in the context of B cell evolution, both ML and NJ methods require setting the unmutated reference germline segments as the outgroup. In contrast, the Bayesian phylogenetic inference program BEAST inherently produces a rooted topology without the need to explicitly specify an outgroup. This strategy, however, completely ignores any prior information regarding known reference germline sequences and potentially worsens estimation of mutation rates and evolutionary distance. While there is the option to specify the germline sequence as a monophyletic taxon in BEAST, how this influences the resulting clonal lineage remains unexplored. We therefore asked how either including or excluding the reference germline gene segments impacted the clonal lineages for each of the aforementioned rooting strategies while keeping all other parameters constant. To answer this, we randomly sampled 50 time-resolved (containing sequences from more than 1 time point) clonal families per cohort for the four different rooting strategies and estimated the Bayesian phylogenetic trees. We assessed how topology and parameters were impacted for the different settings (Figure 3A). We immediately observed stark differences in tree structure between those clonal lineages with or without the germline (Figure 3B). Surprisingly, this not only included differing branch lengths but also the topology of the maximum clade credibility (MCC) tree, as demonstrated by the presence of clades containing altered clonal relationships despite identical sequences between trees (Figure 3B). After visualizing these differences between trees either containing or excluding the reference germline, we decided to quantify how various output parameters from BEAST were impacted across the different rooting strategies. We first calculated how correlated the mean posterior probability was for each tree containing identical sequences excluding the reference germline for those clonal lineages grouped by V- and J-genes plus CDR3 length (Figure 3C). While there was high correlation between the mean posterior probability for all cohorts, cohort specific differences were observed for those trees arising from uninfected versus infected mice (Figure 3C). We finally asked if this trend was consistent across the four different rooting methods. To quantify this, we calculated the log2 ratio for multiple output parameters from BEAST, including the mean posterior probability, substitution rate, tree height and accompanying effective sampling size values (Figures 3D, 3E, S3). This analysis demonstrated effects arising from both infection status and clonal lineage assignment strategy across multiple parameters. The trend that acute and chronic trees had a higher posterior probability, decreased tree height, and increased ESS values when the reference germline was included as the monophyletic group was consistent across rooting strategies (Figure S3). A similar trend was observed when comparing the mean substitution rate for each tree, with the inclusion of the reference germline consistently resulting in a increased substitution rate and decreased tree height for the acutely and chronically infected cohorts (Figure 3D, 3E, S3). While the ESS values for identical MCMC chain lengths for the substitution rate and posterior probability were comparable between trees either with or without the reference germline included, the ESS values for tree height were consistently higher for those trees including the germline sequence (Figure S3). Indeed, the mean tree heights were on average closer to 70 days when the germline reference sequences were set as the monophyletic outgroup, which closely resembles the actual biological sampling and infection scheme (Figure 3E). Together these data show that incorporating germline information in a Bayesian phylogenetic framework alters tree topology and accompanying parameter estimates, in addition to improving tree height estimates.

**Figure 3.**
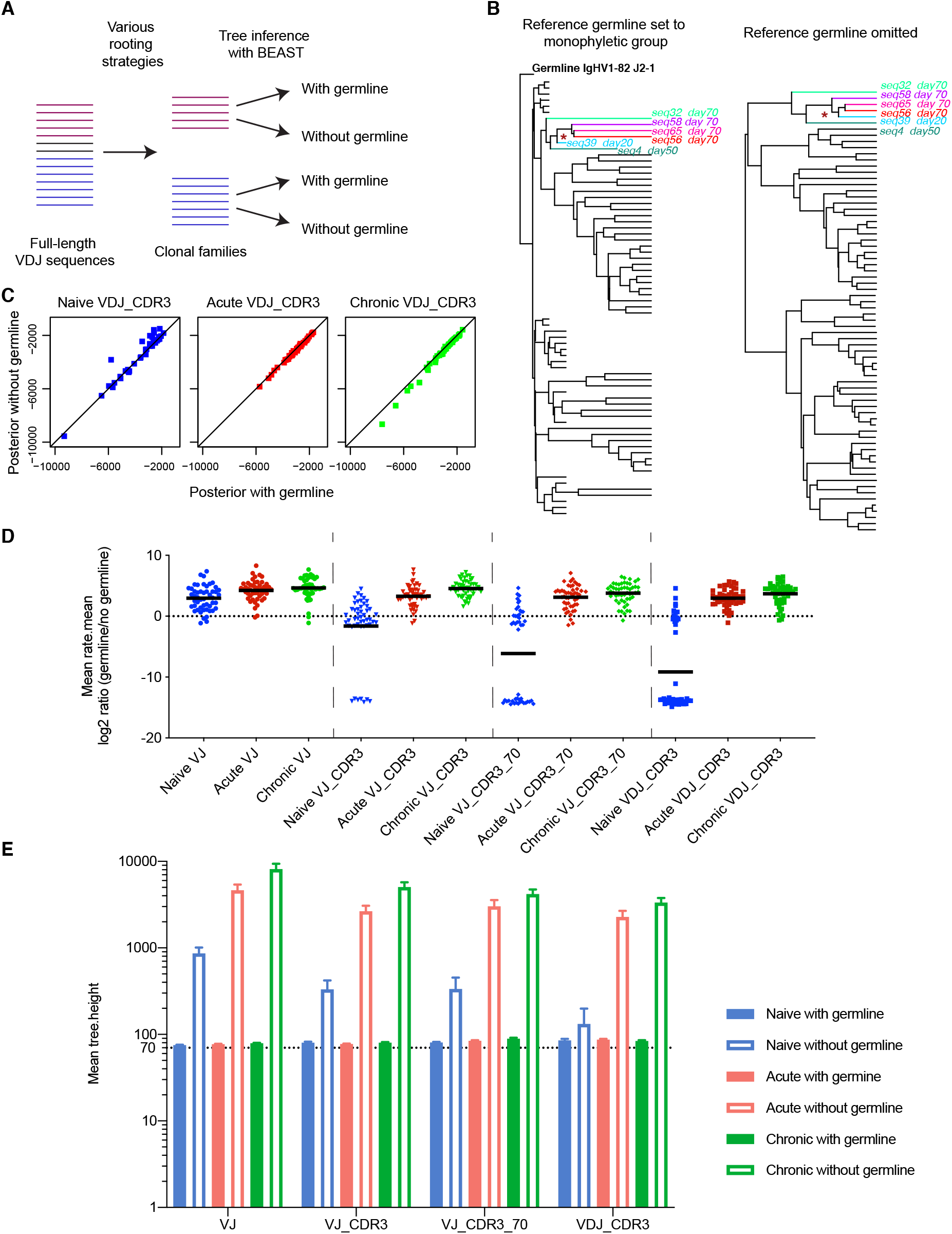
Inclusion of reference germline in Bayesian inference impacts parameter and topology estimates. (A) Schematic depicting Bayesian reconstruction workflow comparing clonal lineages either with or without the reference germline V, (D), and J segments included. (B) Two example phylogenetic trees from the same clonal lineage from a single mouse, either with or without including a reference germline as a monophyletic taxon. Sequences with the same color correspond to identical sequences across trees. Star indicates an example clade that differs between trees due to including the reference germline as monophyletic taxon. (C) Correlation between posterior probabilities for identical clonal lineages either with or without the reference germline as monophyletic outgroup. (D) The log2 ratios of the mean substitution rate estimates from each of the sampled BEAST trees either with or without including the reference germline as the monophyletic outgroup for each cohort and rooting strategy. Color, shape, and line indicate the cohort, rooting strategy, and mean, respectively. (E) Estimated mean tree height for the various rooting conditions. The mean tree height either with or without germline sequence was calculated for all sampled clonal lineages for each cohort across the four indicated rooting methods.

### Different strategies impact the inferred public clonal lineages

While the presence of identical B cell receptor sequences shared across multiple individuals (public clones) has been previously reported, how these public clones evolve across multiple individuals has not been thoroughly studied. Cross-donor phylogenies are a useful approach to infer the evolutionary history from closely related clones across multiple individuals (Zhou *et al*., 2013). This involves performing the aforementioned clonal lineage assignment process on all sequences across all individuals and subsequently inferring the phylogenetic tree using the germline reference as the root. Therefore, we wanted to characterize how the various clonal lineage assignment methods dictated the biological conclusions regarding convergent selection of “public clonal lineages”, or clonal families containing sequences from multiple mice. After clustering sequences and inferring phylogenetic trees, we analyzed how many trees contained sequences from multiple cohorts (Figures 4A, 4B). We immediately observed that grouping sequences sharing V- and J-gene segments resulted with the majority of cross-donor phylogenies containing B cells from all three cohorts (Figure 4B). Indeed, we observed that this proportion of public lineages decreased upon implementing more stringent clonal assignment strategies, with more “cohort-private” lineages arising when requiring identical CDR3 length, sequence homology or identical D-gene alignment. There was higher public lineage overlap between the chronic and acute cohorts, as one might expect due to their similar infection histories. Surprisingly, the rooting strategy influenced the proportion of trees containing either chronically and acutely infected mice or all three cohorts, even between those rooting strategies differing only by a homology requirement (Figure 4B). Similarly, differences between the VJ_CDR3 and VJ_CDR3_70 rooting strategies were observed when comparing the average representation of each mouse in the cross-donor phylogenies (Figure 4C). There was a small proportion of trees containing all mice from the given cohort when grouping by VJ_CDR3, but this was no longer observed after incorporating either a homology threshold or demanding matching D-gene alignments (Figure 4C).

**Figure 4.**
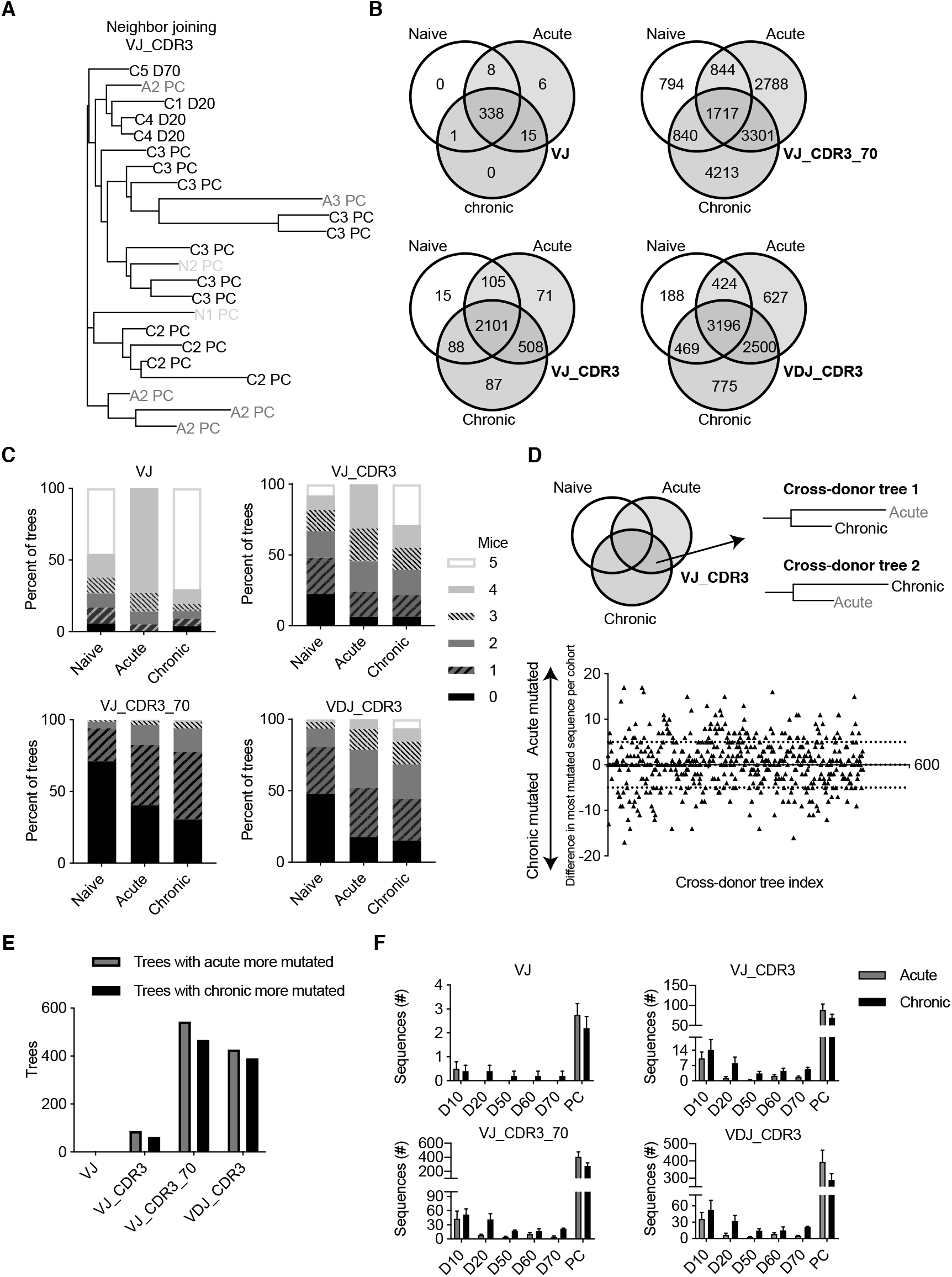
Cross-donor clonal lineages reveal clonal convergence across mice and cohorts. (A) Example cross-donor phylogeny containing sequences from multiple mice and cohorts. Name indicates cohort and mouse (A, C, N for acute, chronic, naïve cohort respectively) and time point post infection. Tip labels with “PC” in name correspond to sequences found in the bone marrow plasma cells (PC) at 70 days post infection. (B) Number of cross-donor phylogenies containing sequences from the indicated cohort under different rooting combinations. (C) Percentage of cross-donor phylogenies containing the indicated number of mice from each cohort. (D) Mutational distance from the reference germline sequence for those cross-donor trees containing sequences from both acutely and chronically infected mice. For each tree, the difference in V gene mutations between the most mutated sequence of either cohort was calculated and plotted for the VJ_CDR3 rooting method. The dotted lines separate those trees where either the most mutated sequence from acute (y>5) or chronic (y<-5) contained more than 5 V gene mutations than the other cohort. (E) The number of cross-donor trees containing sequences from both chronically and acutely infected mice where the most mutated sequence of the indicated cohort contains at least five more substitutions than the most mutated sequence from the remaining infection cohort. (F) Number of sequences from indicated time points for those crossdonor trees containing both chronic and acute sequences.

We next questioned if the four distinct clonal family assignment strategies impacted the mutational burden for the two infected cohorts. To answer this, we first computed the maximum number of mutations from the reference germline sequence for each clonal lineage containing both chronic and acute sequences (Figure 4D). Next, we calculated and plotted the difference of maximum mutational burden between the two cohorts for each of the previously selected cross-donor phylogenies.

Enumerating the number of trees with chronic-dominated or acute-dominated mutational load revealed that the clonal rooting strategy had little impact on the trend that both cohorts underwent similar degrees of somatic hypermutation (Figure 4E). Surprised by this finding, we finally investigated whether differences in temporal-resolution were present in those cross-donor trees. We therefore quantified the proportion of sequences from each time point within the public clonal lineages shared between the acutely and chronically infected cohorts (Figure 4F). Our analysis revealed that the public clonal lineages, on average, contain a higher number of sequences from each time point before the terminal sacrifice (Figure 4F), consistent with previous findings (Kräutler *et al*., 2020).

### Biological sampling influences phylogenetic fingerprint

Our previous analyses revealed that the computational processing pipeline dramatically influences the resulting repertoire phylogenetic snapshot. In our comparison, we employed multiple, time-resolved, biological samples for each mouse, including both blood and bone marrow B cell repertoires. However, peripheral immune organs are rarely available in the case of human antibody repertoire sequencing, prompting the question of how well does the phylogenetic fingerprint from the blood match that of lymphoid tissues. Furthermore, the impact of the rooting strategy may further exacerbate differences arising due to biological sampling limitations. To better characterize the interplay between the rooting strategy and biological sampling, we inferred the evolutionary history for clonal lineages containing sequences from either all blood repertoires for a given mouse (10, 20, 50, 60, 70 dpi), only blood repertoires 70 dpi, or only the PC repertoire 70 dpi (Figure 5A). While both pooling all blood repertoires or including solely PC repertoires again revealed a trend that chronically infected mice had a higher number of clonal lineages despite the same amount of starting biological material, when considering only repertoires at 70 dpi the acutely infected animals actually contained the most clonal lineages per mouse (Figures 5A). The rooting strategy did not, however, largely impact the mouse- and cohort-specific effects excluding the VJ rooting strategy where all three cohorts had similar number of clonal lineages (Figure 5A). When quantifying the size of these clonal lineages, the acute cohort again showed, on average, clonal lineages containing more sequences than the other two cohorts. This effect was not observed in the other two sampling schemes where only PC or pooled blood repertoires were included, possibly due to the lack of LCMV-specific B cells in circulation in the acute and naïve cohorts. Despite these differences in the quantity and size of the clonal lineages per each cohort, more subtle differences were observed when employing metrics that quantify tree imbalance, such as the Colless number and the Sackin index (Figure S4). Taken together, these data indicate potential drawbacks from sampling B cell repertoires at a single time point.

**Figure 5.**
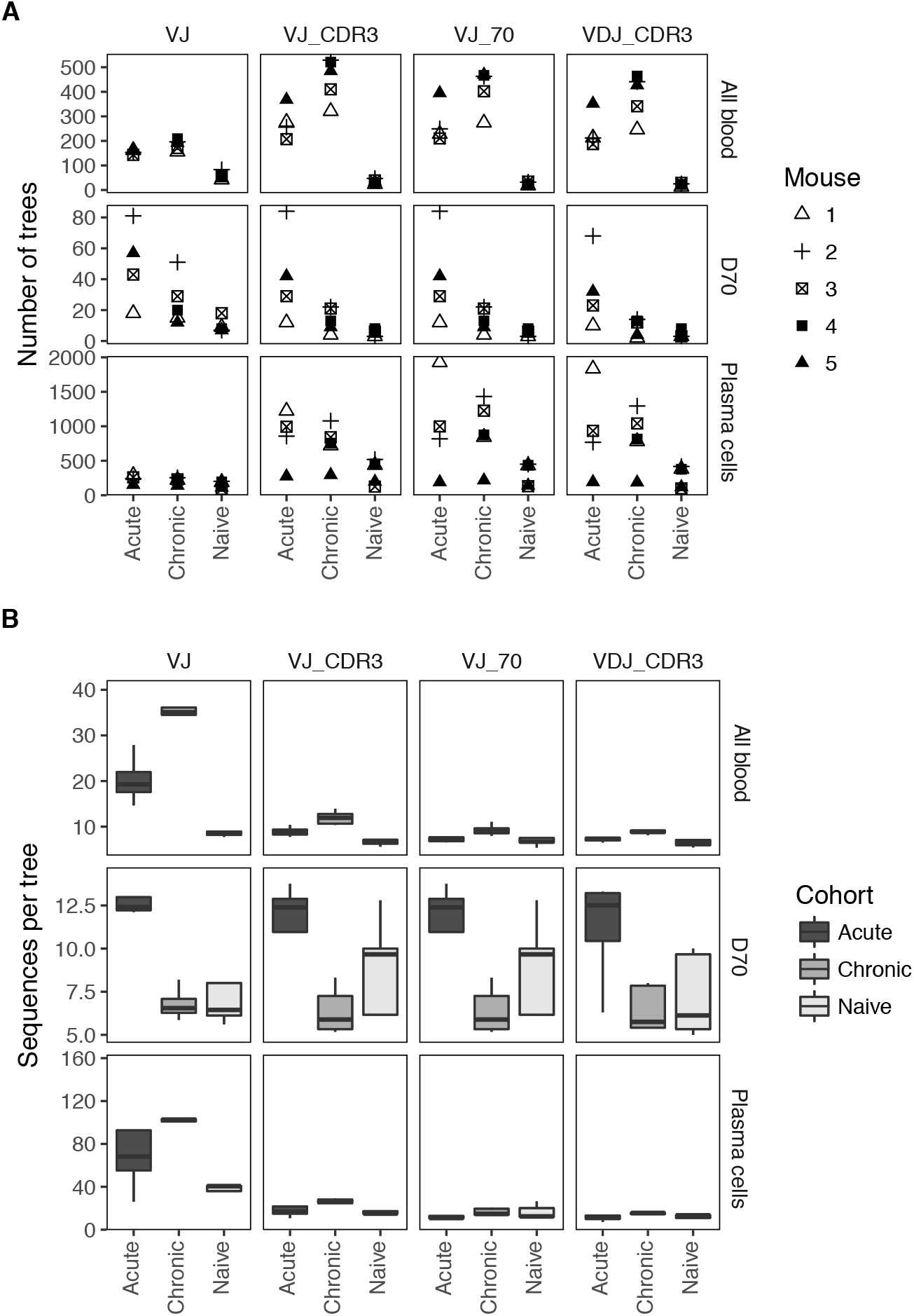
Biological sampling of B cells influences repertoire phylogenetic fingerprint. (A) The number of clonal lineages per mouse and (B) the average number of sequences per clonal lineage obtained from all unique IgG blood-derived sequences from all repertoire at each time point (All blood), all sequences found in blood repertoires 70 days (D70) post infection, or only sequences found in bone marrow PC repertoires 70 days post infection (PCs).

## Discussion

Here we have systematically characterized the influence of the bioinformatics and phylogenetic pipeline on the interpretation of antibody repertoire sequencing data. The strategies discussed here have widely been incorporated in previous studies analyzing B cell data (Stern *et al*., 2014; Tsioris *et al*., 2015; Jackson *et al*., 2014; Heiden *et al*., 2017; Vollmers *et al*., 2013; Wu *et al*., 2015; Zhu, Ofek, *et al*., 2013). It remains unclear, however, just how robust these biological conclusions are to parameter and pipeline variation. We therefore utilized a time-resolved antibody repertoire sequencing data set where B cell repertoires from both blood and lymphoid organs were sequenced for multiple mice across three different cohorts. The three cohorts (uninfected, acutely infected, and chronically infected) represent three conditions in which differing selective pressures would be expected given the importance of B cells in the clearance of chronic LCMV virus (Battegay *et al*., 1993; Hangartner *et al*., 2003). We could thereby use identical B cell sequences as input to various clonal lineage assignment strategies and phylogenetic inference strategies and compare how the mouse- and cohort-specific phylogenetic fingerprint was impacted. Our results stress both the need for caution when interpreting evolutionary conclusions arising from bulk Ig-seq experiments and reveal certain pitfalls to be avoided when conducting such an analysis.

One of the most apparent findings from our analyses is that the clonal lineage assignment strategy can dramatically alter metrics describing the size and number of clonal lineages (Figures 1, 5), the tree topology (Figures 2, 3), the mutational burden (Figures 3, 4) and the degree of convergent selection between cohorts (Figure 4). The various clonal lineage assignment strategies all relied upon an initial alignment to the reference murine germline V-, D- and J-genes, a practice common to many Ig-seq studies. Certain heuristics were additionally applied to some groups, including a CDR3 length requirement due to the biological evidence for minor insertions and deletions during somatic hypermutation (Smith *et al*., 1996) or a certain degree of homology across the CDR3 region. While imposing a restriction upon the CDR3 length has a biological basis and is minorly impacted by parameter fine-tuning, imposing homology requirements upon clonal lineages can vary the biological results compared to simply grouping by V- and J-gene germline alignment and CDR3 length alone. Recently, multiple tools have been specifically developed to perform clonal lineage assignment without the need to set arbitrary thresholds (Ralph and Matsen IV, 2016; Briney *et al*., 2016). The continued validation of such tools on both simulated and experimental data, in addition to increasing the user-accessibility, will hopefully move the field towards a consensus regarding rooting strategies.

The goal of the phylogenetic analysis further dictates how necessary it is to benchmark multiple pipelines. For example, previous work has utilized Ig-seq data and phylogenetics to discover novel HIV-neutralizing antibodies by selecting B cell sequences closely related to a known antibody (Zhu, Ofek, *et al*., 2013). In this particular example, sequences were assigned to the tree if they contained the same V- and J-gene germline elements. This choice of rooting strategy would not necessarily alter which antibodies are closest to the known HIV-neutralizing antibody, and thereby would not have impacted candidate selection. Other work describing the evolution of HIV-specific antibodies grouped B cell sequences to the clonal family of a known neutralizing antibody in a similar method and subsequently calculated the mutation rate using BEAST (Wu *et al*., 2015). Our findings suggest that other methods of clonal lineage assignment would impact this inferred mutation rate and further stress the need for caution when interpreting such a metric.

While the clonal lineage assignment strategy may not directly impact the aforementioned HIV-neutralizing candidate selection, our results suggest that the phylogenetic inference method would have substantial consequences (Figure 2). We observed extremely low convergence of phylogenetic tree topologies inferred using two different methods on identical input sequences (Figures 2A–2C). Increasing the stringency of the rooting strategy resulted in high convergence between the inference methods, presumably because the sequences included in each clonal lineage are necessarily more similar than if one were to only group by V- and J-genes. Consistent with our findings that distinct topologies were produced by the two inference methods, we observed differences in the accompanying metrics quantifying tree structure when comparing clonal lineages with identical sequences (Figure 2D-F). While the Colless number remained highly correlated for all three cohorts when comparing trees produced using identical input sequences, we observed that the variation increased trees with larger normalized balance. This high degree of correlation suggests an irrelevance of the phylogenetic inference method when considering downstream conclusions; other metrics such as the Laplacian spectra were able to clearly separate the two methods, further highlighting the influence the tree inference method holds. It is worth mentioning, however, that our inference pipelines did not incorporate B cell specific substitution models or include clonal frequency information, both of which are crucial biological considerations to the humoral response. Further work validating and comparing the various tools incorporating these two considerations (Yaari *et al*., 2013; Hoehn *et al*., 2017; DeWitt *et al*., 2018) may reveal even more dramatic differences between inference methods. Future studies could additionally investigate how incorporating dependences between clonal lineages within a single repertoire influences the overarching evolutionary landscape (Hoehn *et al*., 2019).

We have additionally demonstrated a potential benefit to including the reference germline as the monophyletic group when estimating tree heights and mutation rates in a Bayesian framework. This benefit seems to be particularly clear for the clonal lineages of acutely and chronically infected mice, indicated by both higher ESS values despite the same chain length of the MCMC algorithm and that the estimated tree heights more closely resembled the span of the experiment of 70 days (Figures 3E, S3). How to exactly use the reference germline as the monophyletic group in calendar time remains relatively unexplored, but has important repercussions for the aforementioned Bayesian analysis of HIV neutralizing antibodies (Wu *et al*., 2015). Here we set the sampling time (tip date in terms of BEAST) monophyletic group to one day after the start of the infection. This assumes that evolution of the given clonal lineage begins upon infection, and implies that finding the initial precursor B cell at a later time point has low probability. This may partially account for why the naïve cohort behaved differently than the other two infection cohorts when the reference germline was included. More analysis is required to understand if incorporating a sliding window root height for the monophyletic group would benefit those clonal lineages that are later recruited into the B cell response or when the onset of clonal evolution is unknown. Furthermore, we have demonstrated that the inclusion of the germline reference sequence in BEAST can alone impact the tree topology, similarly holding important ramifications when incorporating B cell evolution into the Bayesian framework.

Finally, we utilized cross-donor phylogenetic analysis to better characterize the interplay between clonal lineage assignment and convergent evolution. This style of analysis has been previously performed to discover HIV-neutralizing antibodies by creating phylogenetic trees from multiple patients, some of which contained known virus neutralizing antibodies (Zhu, Wu, *et al*., 2013). Our cohort-focused analysis revealed that the number and proportion of public lineages was dramatically influenced by clonal rooting strategy (Figure 4B). We additionally observed that the rooting strategy impacts the proportion of mice from each cohort that were represented in each clonal lineage (Figure 4C); a relevant finding when constructing cross-donor phylogenies for multiple patients exposed to the same pathogen. Those clonal families containing sequences from the most individuals could represent therapeutic candidates that are potentially broadly neutralizing. Therefore, understanding how the bioinformatics pipeline influences this read out if of crucial importance.

Together, our data both demands increased caution when interpreting evolutionary conclusions from Ig-seq experiments and quantifies the influence of the bioinformatics pipeline on biological conclusions. Further scrutinizing how we quantify B cell evolution and selection in response to immunizations and pathogens is crucial for the development of therapeutics that are effective across a wide range of individuals. In conclusion, we demonstrate the benefit of comparing multiple bioinformatics pipelines to ensure robust evolutionary results including antibody forest size and structure, mutation rate, and tree topology,

## Methods

### Repertoire processing

Ig-seq data from accession number E-MTAB-8585 (Kräutler *et al*., 2020) was aligned using MiXCR (v2.1.2) to the built in murine reference germline segments under default parameters (Bolotin *et al*., 2015). The alignments were subsequently clonotyped by the full length VDJ region on the nucleotide level and exported using exportClones function with presets set to full. The clonotypes that successfully aligned to both the IgG constant region and both a V, and J alignment were retained for phylogenetic analysis. In the case of trees arising from the VDJ_CDR3 rooting strategy, only those clones containing a D alignment were retained. Clonal lineage assignment was performed on the V, (D), and J gene segments with the highest alignment score for each clone. Only one unique sequence per full length VDJ clone was included in each root, thereby ignoring clonal abundance information. For those rooting strategies involving homology requirements (VJ_CDR3_70, VJ_CDR3_80, VJ_CDR3_90, VJ_full_70, VJ_full_90), homology was calculated by the edit distance normalized by sequence length (Levenshtein I, 1966) for either the CDR3 region (for VJ_CDR3_70, VJ_CDR3_80, VJ_CDR3_90) or the entire VDJ sequence (for VJ_full_70, VJ_full_90). Only those clonal lineages containing three or more sequences were included in the analysis.

### Phylogenetic inference

For each clonal lineage, the corresponding V, (D), and J reference genes from IMGT were appended together and added to the list of sequences. If there were multiple alleles present for the given V, (D), or J reference gene, the first one was selected (*01 in IMGT annotation).

Neighbor joining trees were inferred by first computing the pairwise edit distance between each sequence using the stringdistmatrix function in the R package stringdist (Van der Loo, 2014). The resulting distance matrices were then used as input to the nj function in ape (Paradis *et al*., 2004). The germline reference sequence was set as the root using the reroot function from the R package phytools (Revell, 2012). For ML phylogenies, a multiple string alignment was performed using clustalw2 under default parameters for each clonal lineage to produce a nexus file. For ML trees, the nexus files were first converted to phylip files using readseq and then used as input to raxml using a GTRgamma *model* and with the reference germline set to outgroup. For Bayesian analysis, additional sub-sampling was performed due to computational demands. 50 clonal lineages containing sequences from at least two time points were randomly sampled from each cohort across the four rooting strategies (VJ, VJ_CDR3, VJ_CDR3_70, VDJ_CDR3). If a clonal lineage exceeded 100 members, 100 randomly sampled sequences were selected. Multiple string alignments were performed as previously mentioned twice for each clonal family, once with the reference germline sequence included and once without. Resulting nexus were first converted to xml files using a custom R script that incorporated the sampling time as the tip date and set the reference germline as the monophyletic group with the tip date set to one. BEAST (version 2.5) was used to infer phylogenetic trees for each of the 50 randomly sampled clonal lineages per cohort. The GTR substitution model was used with gamma-distributed site heterogeneity. Divergence times were estimated using a relaxed lognormal clock model with the clock rate set to 1 and the discrete rate count set to −1. A birth death prior was used with uniformly distributed prior probability distributions. The chain length of the MCMC algorithm was set to 8,000,000 and was logged every 5000 iterations. The MCC tree was extracted using the program TreeAnnotator (Bouckaert *et al*., 2014) with a 10% burn-in as previously described (Yermanos *et al*., 2017). Mean parameter estimates and ESS values were summarized from the output log files using TreeLogAnalyser, part of the BEAST2 suite (Bouckaert *et al*., 2014).

### Topology metrics

After inferring trees for each clonal lineage using the aforementioned inference pipeline (two trees for each clonal lineage, using either ML or NJ), the Robinson-Foulds distance between each pair of lineage trees was calculated using the R package phangorn (Schliep, 2011) and normalized by the number of sequences within in tree. Laplacian spectra were calculated for each pair of lineage trees using the spectR package Rpanda as suggested by the developers (Lewitus and Morlon, 2016). The mean, maximum, minimum, and standard deviation of spectR’s output parameters (principals, asymmetry, peakedness1, peakedness2, and eigengap) were calculated for all clonal lineages within each mouse. This matrix (parameter versus mouse) was used as input to the base R function prcomp to calculate the principal components. Colless number and Sackin index were calculated using the R package Phylotop and normalizing by the number of sequences within each tree.

Cross-donor phylogenies were computed by first pooling all unique VDJ clones for all mice and all time points and then rooted based on the four aforementioned rooting strategies. The maximum distance from reference germline for each cohort was calculated based on the number of amino acid substitutions relative to the MiXCR determined reference germline sequence.

