## Supplementary materials for "The influence of the phylogenetic inference pipeline on murine antibody repertoire sequencing data following viral infection"

Supplementary material for “The influence of the phylogenetic inference pipeline on murine antibody repertoires sequencing data following viral infection”, Yermanos et al.

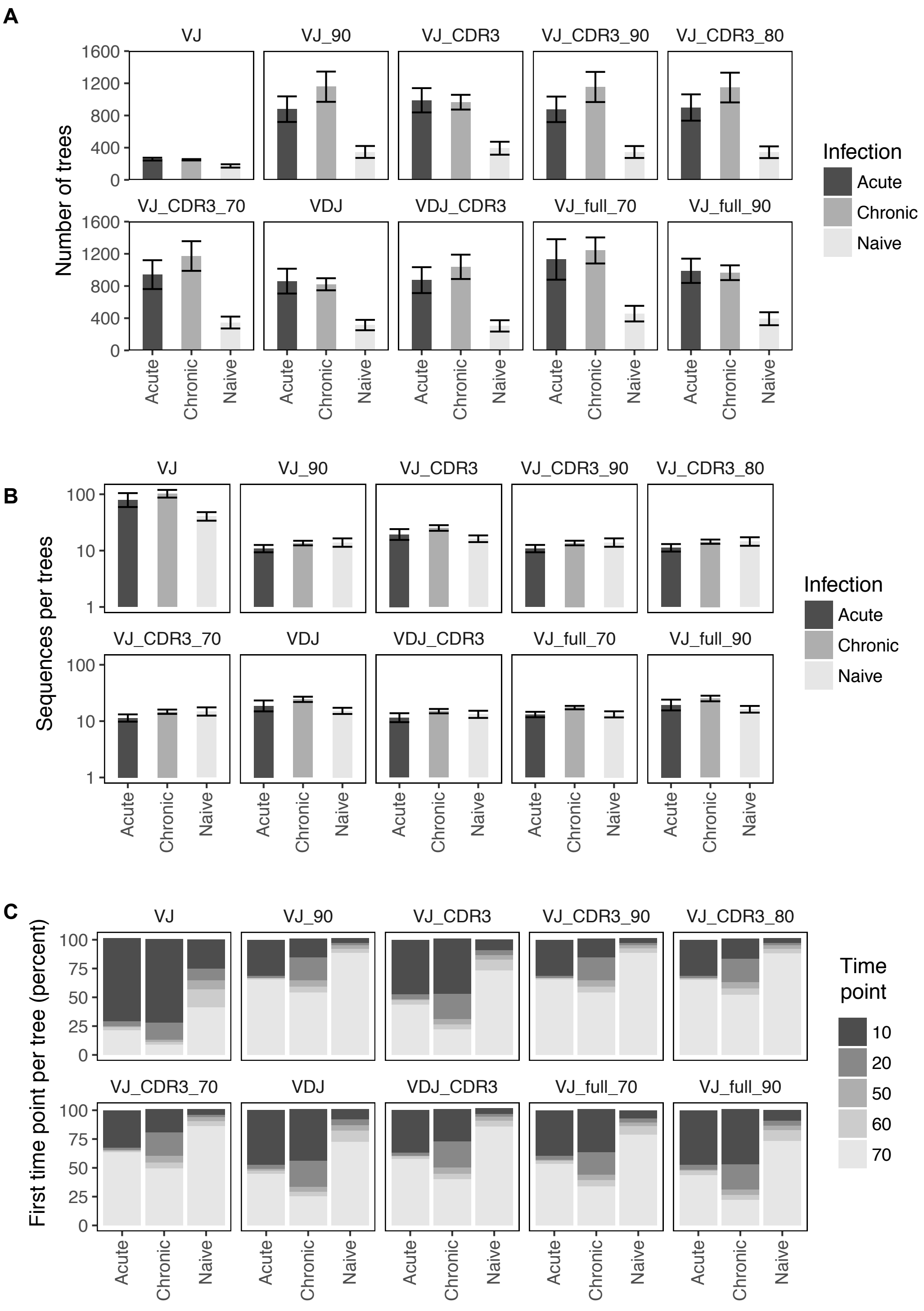


**Figure S1. Size and temporal resolution of clonal lineages across ten rooting strategies.** (A) The mean number of trees for each mouse under various rooting strategies arising from the same alignment. (B) The mean number of sequences within each tree. (C) The average percent of trees within each cohort starting with the indicated time points. Error bars indicate the standard error of mean.

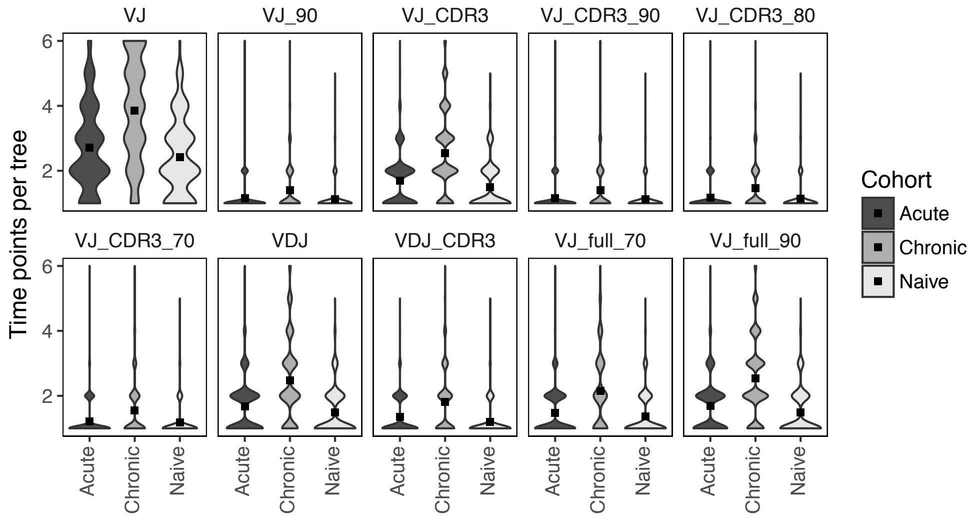


**Figure S2. Distribution of time points found within each clonal lineage.** For each rooting strategy, all clonal lineages from a given cohort were grouped and the number of distinct time points for each tree was calculated. Box indicates the average number of time points sampled across all clonal lineages of the indicated cohort.

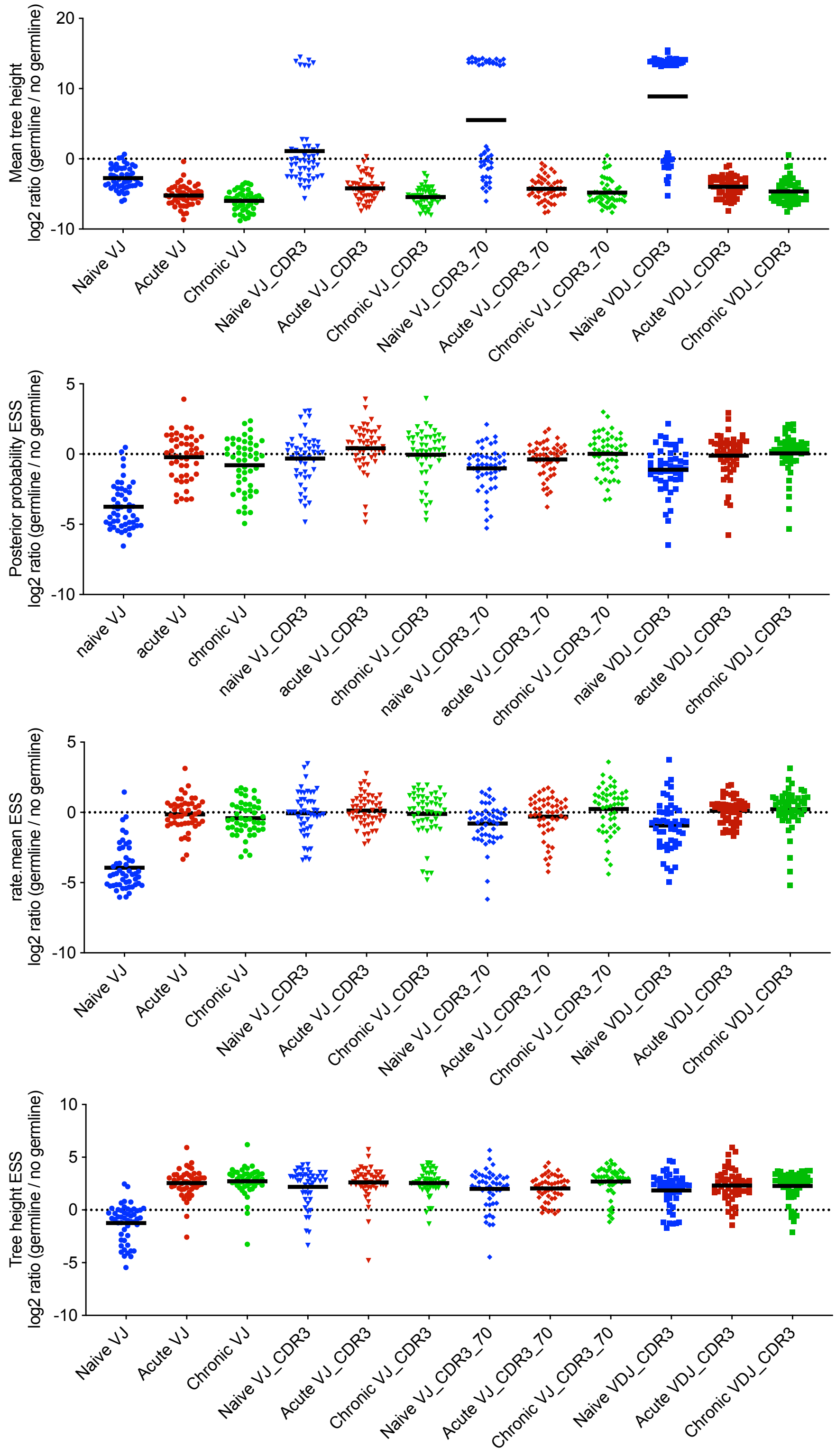


**Figure S3. Inclusion of reference germline segments impacts on tree height and effective size sampling (ESS) values after phylogenetic inference with BEAST.** The $log_2$ ratios of various output parameters from BEAST were compared for clonal lineages containing identical sequences either with or without the inclusion of the reference germline.

**A**

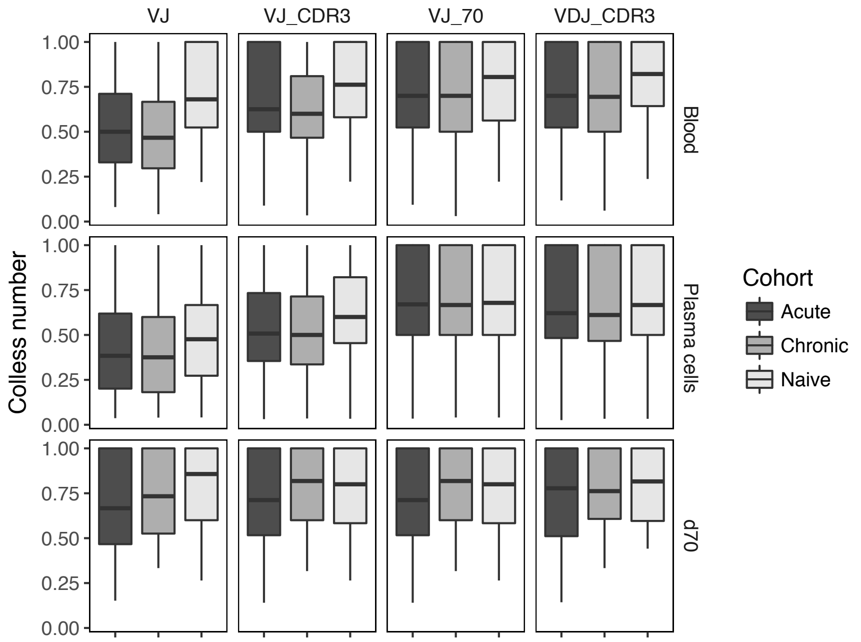


**B**

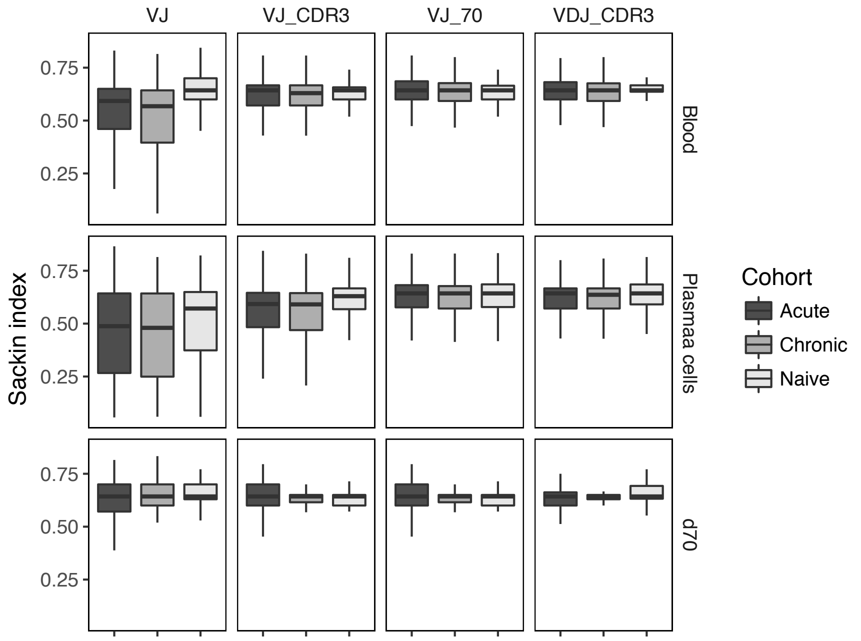


**Figure S4. Biological sampling alters tree imbalance metrics. (A) Normalized Colless number and (B) normalized Sackin index describing how different biological sampling influences tree imbalance.** Clonal lineage assignment was performed on pooled sequences from the indicated time points for each mouse and trees were subsequently inferred with ML. The Sackin index and Colless number were calculated after normalizing for the number of sequences within each tree. Line within each boxplot indicates the mean across all trees of the indicated cohort.
